# Induced defenses alter the strength and direction of natural selection on reproductive traits in common milkweed

**DOI:** 10.1101/093609

**Authors:** Ken A. Thompson, Kaitlin A. Cory, Marc T. J. Johnson

**Author notes:** corresponding author; email—.

## Abstract

Evolutionary biologists have long sought to understand the ecological processes that generate plant reproductive diversity. Recent evidence indicates that constitutive antiherbivore defenses can alter natural selection on reproductive traits, but it is unclear whether induced defenses will have the same effect and whether reduced foliar damage in defended plants is the cause of this pattern. In a factorial field experiment using common milkweed, *Asclepias syriaca*, we induced plant defenses using jasmonic acid (JA) and imposed foliar damage using scissors. We found that JA-induced plants experienced selection for more inflorescences that were smaller in size (fewer flowers), while control plants only experienced a trend toward selection for larger inflorescences (more flowers); all effects were independent of foliar damage. Our results demonstrate that induced defenses can alter both the strength and direction of selection on reproductive traits, and suggest that antiherbivore defenses may promote the evolution of plant reproductive diversity.

---

Understanding the ecological and evolutionary processes that have generated the astounding diversity of plant reproductive traits is an important and longstanding problem in evolutionary biology (Barrett, 2010). Although plant-pollinator interactions have played a major role in this diversification (Stebbins 1970; Harder and Barrett 2007), it is increasingly apparent that both herbivores and antiherbivore defenses have also contributed to the evolution of plant reproductive diversity (Galen, 1999; Irwin *et al.*, 2004; Armbruster *et al.*, 2009; Hanley *et al.*, 2009; Thompson & Johnson, 2016). While we are beginning to understand how herbivores can directly select on reproductive traits (Strauss & Whittall, 2006; Carr & Eubanks, 2014), we know relatively little about the ecological mechanisms underpinning the evolutionary interactions between plant defense and reproduction (Campbell, 2014; Johnson *et al.*, 2015). Here, we test whether induced responses associated with chemical and physical defense traits (hereafter ‘induced defenses’ *sensu* Karban and Baldwin [1997]) alter natural selection on plant reproductive traits.

Antiherbivore defenses may alter natural selection on plant reproductive traits through two non-mutually exclusive mechanisms. First, allocation trade-offs between defense and reproduction may cause investment in defenses to directly reduce reproductive fitness. Such a trade-off could drive selection for increased allocation into reproduction in favour of reduced allocation to defense, and vice versa (Thompson & Johnson, 2016). Numerous studies have shown that induced defenses can change reproductive trait expression (e.g., Hoffmeister et al. 2016) and fitness (e.g., Schuman et al. 2012), but none have tested whether induced defenses alter selection on reproductive traits. Second, because herbivores and pollinators both exert selection on reproductive traits, if defenses alter interactions with either of these selective agents, then phenotypic variation for defense could be associated with patterns of natural selection on reproductive traits (Thompson & Johnson, 2016). While there are theoretical reasons to expect that antiherbivore defenses can alter natural selection on plant reproductive traits (Johnson *et al.*, 2015), there is currently very little empirical evidence to support this hypothesis.

A previous study demonstrated that genetic variation in constitutive defenses (cyanogenesis within *Trifolium repens*) can alter natural selection on reproductive traits (Thompson & Johnson, 2016), but it is unclear whether induced defenses can have similar effects. While both constitutive and induced defenses increase fitness in the presence of herbivores, they differ in that constitutive defenses are always present while induced defenses increase following interactions with herbivores (Karban & Baldwin, 1997). Several studies have shown that, like constitutive defenses, induced defenses can reduce reproductive fitness through allocation trade-offs (Strauss, 1997; Agrawal *et al.*, 1999; Lucas-Barbosa, 2015), and can alter both floral herbivory and pollination (Heil, 2002; Strauss *et al.*, 2002; McCall & Karban, 2006; Kessler *et al.*, 2011). These observations collectively suggest that induced defenses might alter natural selection on reproductive traits.

Despite the similarities between constitutive and induced defenses, they have several important differences that preclude the assertion that both types of defense should similarly alter natural selection on reproductive traits. Constitutive and induced defenses may differ in their associated costs, because constitutive defenses are always expressed, whereas induced defenses are expressed only in response to specific stimuli (Karban & Myers, 1989). Alternatively, induced responses to herbivory can trigger ‘reproductive escape’ whereby plants drastically increase investment into reproduction (Lucas-Barbosa *et al.*, 2013). Constitutive and induced defenses may also have different effects on pollination and herbivory. Induced responses can change floral morphology and scent (Hoffmeister *et al.*, 2016), trigger the release of volatiles that attract parasitoids of herbivores (Schuman *et al.*, 2012; Gols *et al.*, 2015), and have unique effects on herbivore population dynamics and distributions (Underwood & Rausher, 2002; Underwood *et al.*, 2005; Rubin *et al.*, 2015). Because induced and constitutive defenses may differ in their associated costs, and their effects on pollination and herbivory, they may differ in how they influence selection on reproductive traits. Given these differences, experiments that manipulate induced defense are needed to advance our general understanding of how defenses alter natural selection on plant reproductive traits.

When investigating the phenotypic consequences of induced defenses, it is essential to decouple the effects of tissue damage caused by herbivores from the effects of a change in defensive phenotype. This is because herbivory has two main effects on plants: first, it removes tissue that could otherwise be used in photosynthesis; second, it frequently induces chemical and physical defenses. Thus, in natural systems where foliar damage induces defenses it can be difficult to attribute effects to one or the other. With respect to defense-mediated changes in selection, it is especially important to control for foliar damage, because as with defense, foliar damage can affect allocation to reproduction and attractiveness to pollinators (Strauss *et al.*, 1996). The independent effects of leaf tissue removal and induced defenses can be decoupled by applying chemicals to induce the defenses of a plant, and by removing leaf tissue mechanically without inducing defenses. Experimental decoupling of defense and foliar damage is biologically meaningful because species can possess genetic variation for induced responses to a given amount of herbivory (Agrawal *et al.*, 2002), such that the degree to which foliar damage and defense co-vary differs among genotypes. In addition, there are cases where foliar herbivory does not upregulate defenses (Karban & Baldwin, 1997), and where defenses are induced in the absence of damage (e.g., by volatiles; Heil and Karban [2010]). By independently manipulating foliar damage and induced defenses, it becomes possible to quantify their respective contributions to defense-mediated changes in selection on reproductive traits.

Here, we report the results of a field experiment designed to test whether induced plant defenses and simulated foliar damage alter natural selection on reproductive traits in common milkweed, *Asclepias syriaca* L. (Apocynaceae). Our specific research question was: do induced plant defenses alter natural selection on reproductive traits independently of foliar damage? We hypothesized that induced defenses would alter selection on reproductive traits via resource-allocation trade-offs between defense and reproduction, or by altering patterns of pollination and/or herbivory. Evaluating whether induced defenses alter selection on reproductive traits will advance our understanding of the ecological mechanisms driving evolutionary interactions between plant defense and reproduction.

## Materials & Methods

### Study system

Common milkweed, *Asclepias syriaca*, is a perennial plant native to eastern North America (Uva *et al.*, 1997). Plants reproduce vegetatively via rhizomes, and sexually via highly modified flowers (Fig. S1) arranged into umbels (Kephart, 1983). Five nectar-filled hoods surround the fused male and female reproductive organs. The corolla is reflexed and obscures the calyx (Fig. S1). Rather than individual pollen grains, milkweed pollen are aggregated within highly derived structures called pollinia. Pollinia are removed in pairs from flowers as they attach to insect legs and tongues.

Milkweeds *(Asclepias* spp.) are well known for their defensive mechanisms. The common name, milkweed, derives from the thick white latex that exudes from laticifers following mechanical damage to tissues, and it functions by physically impeding herbivores (Dussourd & Eisner, 1987; Agrawal & Konno, 2009). Milkweeds also contain cardenolides, which disrupt sodium-potassium regulation in cells and are toxic to most animals (Malcolm, 1991; Dobler *et al.*, 2012; Zhen *et al.*, 2012). Production of these defenses is upregulated following attack by herbivores, which subsequently reduces herbivory (Van Zandt & Agrawal, 2004). Induction of cardenolides and latex in milkweeds is regulated by the steroid hormone, jasmonic acid (hereafter ‘JA’) (Agrawal *et al.*, 2012), and exogenous JA application upregulates latex and cardenolide production (Rasmann *et al.*, 2009; Agrawal *et al.*, 2012). Milkweed defense traits exhibit heritable variation within populations (Agrawal, 2004), are subject to selection by specialist herbivores (Agrawal, 2005b), and have diverged among species in the genus (Agrawal & Fishbein, 2008; Agrawal *et al.*, 2009). We refer to induction of cardenolides and latex following herbivory or JA application as an induced ‘defense’ because of the substantial evidence that these traits are involved in defense against herbivores.

### Experimental design

To determine whether induced defenses alter selection on plant reproductive traits independently of foliar damage, we manipulated both induced defenses and foliar damage in a 2 × 6 fully factorial field experiment (i.e., 12 treatment combinations) using 156 naturally-occurring milkweed plants on and around (study location hidden during blind review) (approx. 1.25 km^2^; [-] N, [-] W; [-] m a.s.l.). We only included plants with no herbivore damage, and neighbouring plants were separated by at least 1.5 m to reduce the likelihood of sampling the same genet twice (La Rosa, 2015). We covered all plants during the second week of June 2014 with spun polyester bags (Pro 17, Agrofabric, Alpharetta, GA, USA), and tied the bags at the top and bottom of each plant to prevent access by insects (Fig. S1). As plants matured and formed flower buds, we tied the upper-portion of bags below the lowest buds (Fig. S1); this allowed for flower visitation while protecting vegetative parts from herbivores.

We experimentally manipulated induced plant defenses using a treatment with two levels (the ‘induction’ treatment). Plants were either assigned to a control group with no induction (‘control’ plants), or to a treatment group where we used the exogenous application of JA to induce defenses (‘JA-induced’ plants). We prepared a 0.5 mM JA solution by dissolving 100 mg JA (Sigma-Aldrich #77026-92-7) in 2mL of 99.5% acetone and diluting with 950 mL of distilled water; we also prepared a control solution that lacked dissolved JA but was otherwise identical to the treatment solution (Thaler *et al.*, 1996). This method has been used previously to induce defenses in *A. syriaca* (e.g., Mooney *et al.*, 2008). We applied either the JA or control solution to each plant by spraying every fully expanded leaf once (ca. 1 mL leaf^-1^) during each of three applications spaced at two week intervals (13 & 27 June, 11 July); the first application occurred while plants were still undergoing vegetative growth and before any plants had flowered. At the time of initial treatment application, mean plant height was 65.4 ± 14.4 [SD] cm and mean leaf number was 11.49 ± 2.43. We used intervals of two weeks to apply treatments because JA-induced chemical changes are maintained in other systems for at least 14 days (Thaler, 1999).

To decouple the effects of chemically induced defenses and foliar damage, we imposed a leaf-tissue removal treatment (hereafter the ‘damage’ treatment) that had six levels. We imposed our treatment by cutting each leaf on a plant with scissors to remove 0, 10, 20, 30, 40, or 50% of leaf tissue, which encompasses the natural range of damage that *A. syriaca* plants typically experience (Turcotte *et al.*, 2014). We imposed the damage treatment with multiple levels so we could detect any linear or non-linear effects of leaf removal on selection. Leaves were cut once at the start of the experiment, concurrently with the initial application of the induction treatment. We used scissors rather than herbivores because we explicitly wanted to determine the effects of defoliation in the absence of induced defenses; clipping with scissors should not have caused an induced response because of an absence of herbivore saliva, which frequently elicits JA (Agrawal *et al.*, 1999; Musser *et al.*, 2002; Rodriguez-Saona *et al.*, 2002). This rationale is supported by data from Mooney *et al.* (2008), who found that exogenous JA-application—but not mechanical foliar damage—up-regulated the production of induced defenses in *A. syriaca* in ways that were similar to the effects of monarch (*Danaus plexippus*) caterpillar herbivory.

We measured several traits during our experiment. During initial treatment application, we measured plant height, and counted leaves and inflorescence buds. We recorded the first date of flowering for each plant, and measured several aspects of flower size: hood length & width, petal length & width, and the length and width of entire flowers (see Fig. S1). We measured these traits to the nearest 0.5 mm on up to five flowers from the lowest three inflorescences, or all inflorescences for plants with ≤ 3 inflorescences. To obtain mean values of these measurements for individual plants, we averaged all flower size measurements within inflorescences, and then across inflorescences. We counted the number of flowers within the inflorescences used for floral measurements, and refer to the mean number of flowers within inflorescences hereafter as ‘inflorescence size’. In addition, we recorded the total number of inflorescences that each plant initiated, hereafter referred to as ‘inflorescence number’. We harvested the aboveground vegetative biomass of each plant into paper bags at the end of the experiment. After drying plants for 72 h at 50 °C, we removed leaves and weighed the stem tissue as a measure of aboveground biomass. Leaves were removed to avoid including direct treatment effects (mass lost due to leaf tissue removal), or confounded variation in leaf abscission between treatments.

We recorded estimates of both male and female fitness. For male fitness, we counted the number of pollinia removed from five flowers from the lowest three inflorescences, or all inflorescences for plants with ≤ 3 inflorescences. Pollinia removal can be accurately determined on flowers over their entire lifetime, including on senescing flowers. We took the mean number of pollinia removed from each inflorescence, and averaged that among inflorescences to estimate male fitness. Two studies have quantified the relationship between pollinia removal and seeds sired in milkweeds using genetic markers, and found them to be significantly correlated (*r*_Pearson_ range: 0.44-0.47) (Broyles & Wyatt, 1990; La Rosa, 2015). These studies indicate that a large fraction of pollinia removed by pollinators do not sire seeds. Thus, our measure of male fitness represents the upper limit of an individual’s siring success; we acknowledge that this metric is an imperfect estimate of true male fitness, and a more accurate estimate would require molecular markers and exhaustive sampling of the population. As an estimate of female fitness, we harvested ripe fruits and counted the number of viable seeds. We did not include measures of fitness related to vegetative reproduction because *A. syriaca* is a long-lived plant, and it is difficult to determine the belowground rhizome connections between ramets. Nevertheless, this did not affect our ability to effectively address our research questions about whether induced defenses and herbivory alter selection on reproductive traits.

### Statistical analysis

Before analysis, we assessed whether our data met assumptions of normality and homogeneity of variance. We transformed some traits to meet the assumptions of our analyses (Table S1). We calculated pairwise correlations between all measured traits to test for multicollinearity. The six flower size traits were positively correlated (mean *r*_Pearson_ = 0.62), so we collapsed them into two principal components subject to “varimax” rotation using the R package ‘psych’ (Revelle, 2015). The first principal component (PC1) was highly positively correlated with the three flower width measurements (hereafter *floral width PC*), and the other principal component (PC2) was positively correlated with the three flower length measurements (hereafter *floral length PC*); PC1 and PC2 explained 68% and 14% of variation in the data, respectively (Table S2). All pairwise correlations between remaining traits were < 0.7 after collapsing floral measurements except between inflorescence number and biomass (*r*_Pearson_ = 0.78); we retained both variables in spite of this correlation because of an *a priori* interest in analysing differences in defense-mediated changes in selection on both of these traits (Table S3, S4). We also calculated coefficients of phenotypic variation as CV_p_ = 100(V_p_^0.5^ / *μ_i_*), where *μ_i_* is the untransformed mean of trait *i* (Houle, 1992). For all subsequent analyses we standardized traits to a mean of 0 and a standard deviation of 1.

We also used univariate ANOVA to test whether the induction and damage treatments affected the expression of phenotypic traits and fitness components. This allowed us to determine whether there were any allocation costs associated with defense, and to evaluate the phenotypic effects of leaf damage. If damage or induced defenses directly change the expression of a reproductive trait, this could alter selection on the trait.

To determine if JA-induction altered natural selection on plant reproductive traits independently of foliar damage, we used multivariate phenotypic selection analyses (Lande & Arnold, 1983). We first calculated relative fitness (*w_i_*) across all plants, standardized trait values, then ran a linear model regressing relative fitness values against the main effects of each treatment (induction and damage), each phenotypic trait, trait × induction interactions, and trait × damage interactions. Induction was coded as a categorical variable with two levels, and damage was a continuous variable with six levels. The model took the following form:

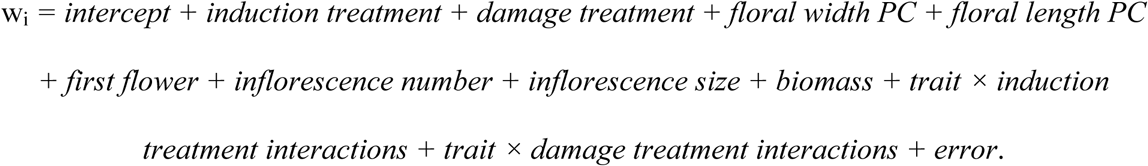

Three-way trait × induction treatment × damage treatment terms and non-linear selection coefficients were included in preliminary models, but all were nonsignificant (*P* > 0.5) and are not reported. Significant main effects for treatments indicate that the treatment influenced fitness, and significant main effects for traits indicate that the trait was under directional selection. A significant trait × treatment interaction indicates that the treatment altered directional selection on the trait (i.e., the confidence intervals on selection gradients for different treatment levels are non-overlapping).

We analyzed multivariate selection models using a multi-model selection procedure implemented using the *dredge* function in the R package ‘MuMIn’ (Barton, 2015). This procedure tests models that span all possible combinations of variables (except for models containing interactions without the corresponding main effects). We took weighted-averages from all models within 2 ΔAIC of the best model—where better fitting models are weighted more heavily—to determine final model-averaged parameters. The best model had 12 variables, and statistically equivalent models (within 2 ΔAIC) had between 7 and 14 variables. We interpreted the ‘full’ model output from the analysis because parameter estimates in the ‘conditional’ output are biased away from zero; thus, the ‘full’ model estimates are more conservative (Barton, 2015). We note that the general conclusions of the present study do not change when the multivariate selection model is analyzed using standard multiple regression (see R script).

Because induced defenses altered selection on traits (see **Results**), we quantified patterns of selection on JA-induced and control plants separately. We restrict these analyses to traits where we detected significant trait × induction treatment interactions. To accomplish this, we analyzed linear models that included all terms from the dredge model with the lowest AIC (i.e., the best fitting individual model; see Table 1) and extracted residuals for fitness and trait values using the approach for partial variance in multiple regression analyses as explained by SAS (SAS Institute Inc, 2009). Specifically, we extracted fitness residuals from the model after removing both the focal trait and the focal trait × induction treatment interaction term. We then extracted trait residuals from an identical model with the standardized trait as the response variable rather than relative fitness. Regression of fitness residuals on trait residuals estimates the selection gradient on the focal trait while controlling for other measured traits. We partitioned the data across treatments, and then analyzed linear models of fitness residuals on trait residuals to estimate the selection gradient in each treatment separately. All data were analyzed using R v.3.0.3. (R Core Team, 2014), and all R-code used to generate results and figures is available online.

**Table 1.** Multivariate phenotypic selection analysis depicting model-averaged coefficients (*β*), *z*-values, and corresponding *P*-values from the ‘dredge’ multi-model selection procedure. Significant coefficients (*P* < 0.05) and interaction terms are highlighted in bold. The model includes all plants that had data collected for all measured traits (*n* = 94 plants). Significant main effects indicate that the treatment or trait directly affected fitness, and significant trait × defense interaction terms indicate that selection on the trait is different between JA-induced and control plants. NA values indicate that a term was not sufficiently important to be included in any model within 2 ΔAIC of the best-fitting model.

|  | Female fitness** |  |  | Male fitness*** |  |  |
| --- | --- | --- | --- | --- | --- | --- |
| | $\beta$ | $z$ | $P$ | $\beta$ | $z$ | $P$ |
| Induction* | **-0.513** | **2.519** | **0.012** | 0.068 | 0.589 | 0.556 |
| Damage* | -0.132 | 0.954 | 0.340 | 0.049 | 0.513 | 0.608 |
| Floral width PC* | -0.138 | 0.937 | 0.349 | -0.178 | 0.868 | 0.385 |
| Floral length PC | 0.001 | 0.036 | 0.971 | 0.103 | 0.797 | 0.425 |
| First flower * | -0.090 | 0.784 | 0.433 | -0.002 | 0.086 | 0.931 |
| Inflorescence number* | -0.210 | 1.090 | 0.276 | 0.090 | 0.634 | 0.526 |
| Inflorescence size* | **0.319** | **1.985** | **0.047** | 0.251 | 1.278 | 0.201 |
| Biomass* | **0.574** | **3.638** | **< 0.001** | 0.000 | 0.004 | 0.997 |
| Induction × Damage* | 0.237 | 1.215 | 0.224 | | NA | |
| Floral length PC × Induction | | NA | | 0.121 | 0.761 | 0.447 |
| Floral width PC × Induction* | 0.202 | 1.283 | 0.200 | 0.045 | 0.429 | 0.668 |
| First flower × Induction | -0.003 | 0.096 | 0.923 | | NA | |
| Inflorescence number × Induction* | **0.856** | **3.239** | **0.001** | -0.039 | 0.320 | 0.749 |
| Inflorescence size × Induction* | **-0.563** | **2.849** | **0.004** | -0.112 | 0.636 | 0.525 |
| Biomass × Induction* | **-0.438** | **2.137** | **0.033** | | NA | |
| Floral length PC × Damage | | NA | | 0.002 | 0.058 | 0.954 |
| Floral width PC × Damage | 0.019 | 0.257 | 0.797 | 0.358 | 1.766 | 0.0774 |
| First flower × Damage | -0.006 | 0.129 | 0.898 | | NA | |
| Inflorescence number × Damage | 0.044 | 0.372 | 0.710 | | NA | |
| Inflorescence size × Damage | 0.010 | 0.183 | 0.855 | -0.026 | 0.298 | 0.766 |
| Biomass × Damage | 0.015 | 0.213 | 0.832 | | NA | |

\*term present in model with lowest AIC (i.e., best fitting model) in *dredge* analysis and used to generate residual values for direct selection analysis (female fitness only). \*\*Number of seeds produced. \*\*\*Mean pollinia removed per flower.

## Results

### Treatment effects on traits and fitness

We did not detect direct effects of the induction treatment or the damage treatment on trait values or fitness in our univariate analyses (Table S5). There were nearly significant effects of the induction treatment on inflorescence size (19% larger in JA-induced plants; ***P*** = 0.07) and biomass (10% smaller in JA-induced plants; ***P*** = 0.10). In our multivariate selection model, which accounted for covariance among traits, we detected a significant effect of JA-induction on fitness, where JA-induced plants produced 20% fewer seeds than control plants (*β* = −0.51 ± 0.20 [SEM], ***P*** = 0.012) (Table 1). We did not detect an effect of damage on fitness in our multivariate analysis.

### Population-wide patterns of natural selection

We observed natural selection through female fitness, but not male fitness, in our multivariate selection analyses (Table 1). At the population level, there was significant directional selection for greater inflorescence size (*β* = 0.32 ± 0.16, *P* = 0.047) and biomass (*β* = 0.57 ± 0.16, *P* < 0.001) via female fitness. We found no evidence that selection was operating via male fitness on any trait main effect or trait × treatment interaction (Table 1), despite large variation in pollinia removal (CV_p_ = 98.8; Table S1).

### Defense-mediated changes in natural selection

Phenotypic selection analyses revealed that the induction treatment altered the strength and direction of natural selection on plant reproductive traits via female fitness (Table 1; Fig. 1). Specifically, JA-induced plants experienced selection for more inflorescences (*β*_JA-induced—inflorescence number_ = 2.81 ± 0.98, ***P*** = 0.003), whereas selection in control plants was non-significant (*β*_control—inflorescence number_ = −0.60 ± 0.58, ***P*** = 0.312); the slopes were significantly different between the two treatments (inflorescence number × induction: ***P*** = 0.001). JA-induced plants also experienced selection for smaller inflorescences (*β*_JA-induced—inflorescence size_ = −1.19 ± 0.48, ***P*** = 0.017), while control plants experienced a trend toward opposing selection for larger inflorescences (*β*_control—inflorescence size_ = 0.74 ± 0.38, ***P*** = 0.056); these slopes also significantly differed between treatments (inflorescence size × induction: ***P*** = 0.004). Finally, we observed significant positive selection on biomass in control plants (*β*_control—biomass_ = 1.47 ± 0.42, ***P*** < 0.001), but found no evidence for selection on biomass in JA-induced plants (*β*_JA-induced—biomass_ = 0.17 ± 0.37, ***P*** = 0.645); as above, these slopes were significantly different between treatments (biomass × induction: ***P*** = 0.033) (Fig. S1). No trait × damage treatment interactions were significant.

**Fig. 1.**
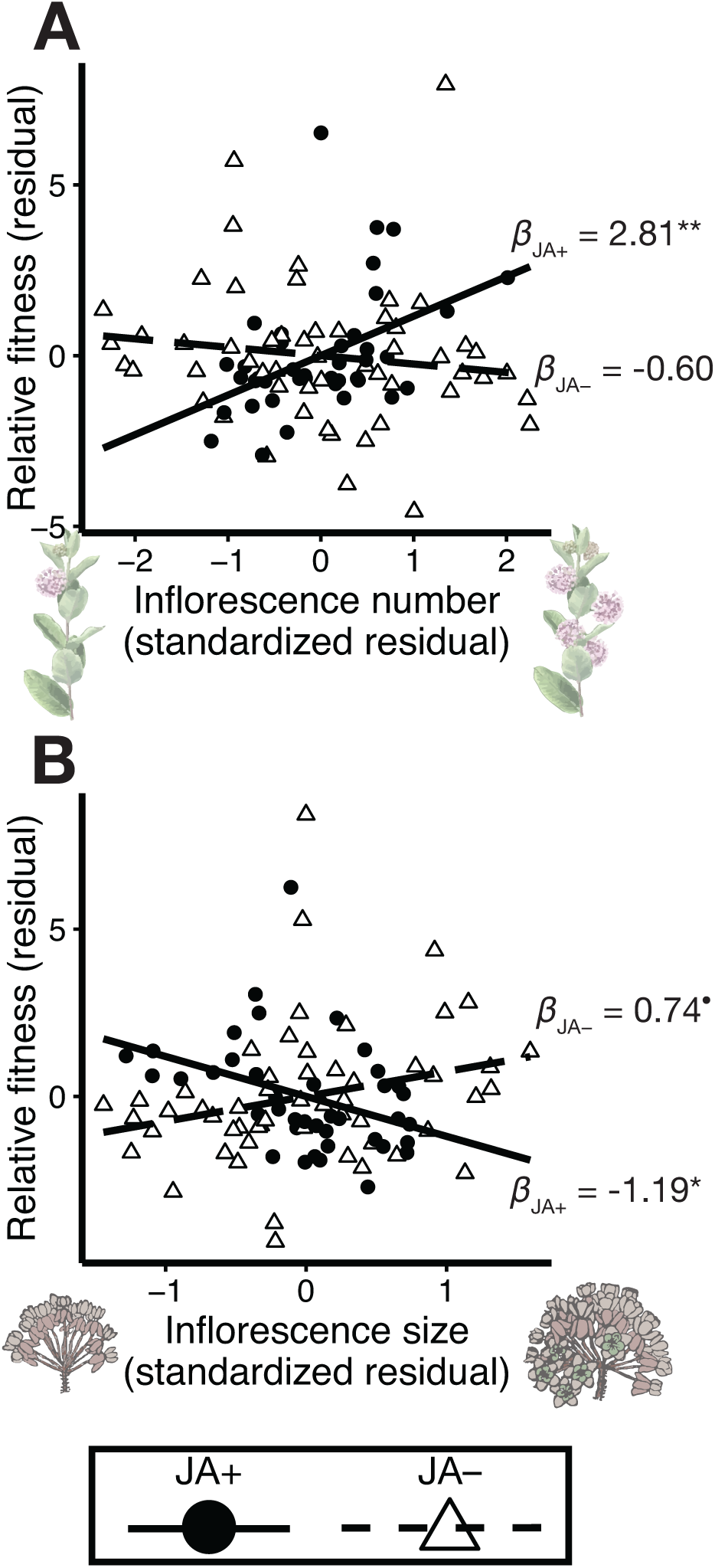
Scatterplots comparing linear selection gradients between JA-induced (*n* = 40) and control (*n* = 54) plants in our experiment. **(A)** JA-induced plants experienced strong positive selection for more inflorescences, but selection was not acting on control plants. **(B)** Plants in both treatments experienced selection on inflorescence size, but selection was negative in JA– induced plants (JA+) and trended toward positive in control (JA−) plants. Selection gradient (*β*) values are reported adjacent to the corresponding selection gradient regression lines (˙*P* = 0.056; \**P* < 0.05; \*\**P* < 0.01). Residual values from regressions with all traits are plotted to visualize the direct effect of the focal trait on fitness while controlling for the effects of other traits in the model.

## Discussion

Studies seldom consider antiherbivore defenses when investigating the evolution of plant reproductive diversity. Recent research, however, highlights independent roles for both herbivory and antiherbivore defenses in shaping natural selection on plant reproductive traits (Campbell, 2014; Carr & Eubanks, 2014; Johnson *et al.*, 2015; Thompson & Johnson, 2016). In this study, we asked whether induced plant defenses alter natural selection on reproductive traits independently of foliar damage. Our results demonstrate that selection on both inflorescence size and number changed following the chemical induction of plant defenses, whereas manual defoliation did not alter selection on any trait. Taken together, our results suggest that plastic phenotypic variation in defense can alter both the strength (coefficient magnitude) and direction (coefficient sign) of natural selection on plant reproductive traits.

### Causes of defense-mediated changes in natural selection

Resource allocation trade-offs between defense and reproduction, where fitness is reduced in highly defended plants, may cause selection for increased allocation to reproduction. JA-induced plants produced fewer seeds than control plants in our experiment, suggesting that there was a direct fitness cost of inducing defenses in *A. syriaca* (Strauss *et al.* 2002). In general, reproduction through female function is more resource-limited than through male function (Haig & Westoby, 1988), and resource limitation is thought to be the most important determinant of seed production in milkweeds (Wyatt & Broyles, 1994). We found that JA-induced plants experienced selection for more inflorescences, but we did not detect selection on inflorescence number in control plants. This may represent selection for higher investment into reproduction in JA-induced plants (Herms & Mattson, 1992). Milkweeds can mature few fruits per inflorescence, and large inflorescences may be especially inefficient in JA-induced plants if many potential fruits are aborted because of insufficient resources. The observed selection for more inflorescences that are smaller in size among JA-induced plants could thus be selection to partition flower production among more inflorescences in order to increase fruit set per inflorescence. Simultaneous manipulation of defense and resource environment is required to test these hypotheses.

In addition to resource allocation trade-offs, altered interactions with agents of selection on plant reproductive traits could cause the observed defense-mediated changes in natural selection. A previous study in *A. syriaca* found that herbivores destroyed > 90% of the smallest inflorescences but negligibly damaged large inflorescences (Willson & Rathcke, 1974). Thus, large inflorescences may allow milkweed plants to tolerate herbivory (Strauss & Agrawal, 1999). Our data indicates that JA-induced plants experienced selection for smaller inflorescences, while control plants experienced a trend toward selection for larger inflorescences. Induced defenses can reduce florivory (McCall & Karban, 2006), and thus selection for larger inflorescences in (undefended) control plants may be selection for tolerance to herbivore damage. Experimental manipulation of both florivory and defense are needed to test this hypothesis.

### Novel contributions of this study

Our study is the second to demonstrate that antiherbivore defenses can alter natural selection on plant reproductive traits. Thompson and Johnson (2016) showed that the cyanogenesis polymorphism of *Trifolium repens* alters natural selection on petal and inflorescence size. The present study builds on this result in two key ways. First, while the previous study manipulated genetic variation for a constitutive defense, we manipulated induced defenses. That our results are qualitatively similar indicates that both constitutive and induced defense, and both genetic and plastic variation for defense, can alter natural selection on reproductive traits. Second, we experimentally manipulated foliar damage, while Thompson and Johnson (2016) measured natural herbivore damage and included it as a covariate. Defense and herbivory were significantly correlated in their study, and thus our study provides a stronger test of the hypothesis that defense-mediated changes in selection on reproductive traits are independent of foliar damage. Thus, our study builds on that of Thompson and Johnson (2016) and provides additional evidence in support of the hypothesis that plant defenses can alter natural selection on reproductive traits.

Common milkweed, *Asclepias syriaca*, is a model plant system for studying the evolutionary ecology of induced responses to herbivory (Malcolm & Zalucki, 1996). Identifying costs associated with induced responses is a major goal in research on induced responses (Agrawal, 2005a), and our study provides the first evidence of fitness costs associated with induced responses in milkweeds. Our results generate testable hypotheses aimed at addressing the ecological mechanisms underlying costs—and ecological consequences—of induced responses in this system. Testing these hypotheses will advance our understanding of the evolutionary ecology of induced responses by furthering the development of milkweeds as a model system for the study of plant-insect interactions.

### Caveats

We acknowledge several limitations of our experiment and analyses. First, the most accurate estimates of selection utilize breeding values, rather than phenotypic variation among individuals, because environmental covariance between traits and fitness can bias estimates of selection in the field (Rausher, 1992; Stinchcombe *et al.*, 2002). While our use of phenotypic data could bias overall estimates of selection, we applied our treatments randomly and it is unlikely that environmental effects would influence our conclusion that induced defenses alter selection. Second, *A. syriaca* is a highly clonal perennial, and our estimates of fitness likely capture a small portion of lifetime fitness. However, our methods still allow us to accurately quantify defense-mediated changes in selection within a generation. Third, although we found no evidence that natural selection was operating through male fitness, our male fitness metric may be too coarse to detect more subtle patterns. Fourth, we did not verify differences in defensive properties across JA-induced and control plants; other studies have confirmed the efficacy of identical treatments in *A. syriaca* (e.g., Mooney *et al.*, 2008), and the strong effects of induction treatment on fitness and selection further indicate that our treatments altered plant physiology. Last, manual defoliation did not affect traits or fitness, which suggests that our damage treatment was not severe enough to impose fitness costs on *A. syriaca.* We note that our result is consistent with other studies demonstrating low fitness consequences of defoliation in *A. syriaca* (e.g., Hochwender *et al.*, 2000), and that higher levels of damage would have been biologically unrealistic for *A. syriaca* (mean % leaf herbivory in the field = 4.0 ± 1.5%, min = 0%, max = 24.2%; Turcotte et al. [2014]). Despite these caveats, our field experiment provides strong evidence that induced defences can alter natural selection on reproductive traits via female fitness in a natural milkweed population.

### Conclusions

Understanding the factors that underlie the diversification of plant reproductive structures is an important area of research in evolutionary biology (Barrett, 2010). Recent evidence demonstrates that both herbivores and antiherbivore defenses promote plant reproductive diversity on both macroevolutionary (Armbruster, 1997; Armbruster *et al.*, 2009; Hanley *et al.*, 2009), and microevolutionary timescales (Thompson & Johnson, 2016). Our results demonstrate that induced plant defenses can alter both the strength and direction of natural selection on reproductive traits independently of foliar damage. While the available evidence suggests that antiherbivore defenses can alter natural selection on plant reproductive traits, very little is known about the mechanisms responsible for these patterns. It is also currently uncertain if disruptive selection associated with defense can ultimately drive phenotypic divergence within populations. Experimental studies that manipulate defense and resource limitation, and defense and pollinators and/or herbivores, are needed to fully understand the ecological mechanisms through which antiherbivore defenses alter natural selection on plant reproductive traits.

## Acknowledgements

The authors are grateful to D. Anstett, D. Carmona, S. Heard, R. Rivkin, J. Santangelo, J. Stinchcombe, & A. Winn for comments that improved the manuscript. KAT was supported by an NSERC CGS-M award and a Queen Elizabeth II Scholarship in Science & Technology. KC was supported by an NSERC USRA. The project was supported by an NSERC Discovery Grant to MTJJ.

## Author contributions

MTJJ and KAC conceived and designed the experiment. KAC collected the data. KAT and MTJJ analyzed the data. KAT wrote the first draft of the manuscript, and KAT and MTJJ revised the manuscript.

## Supplementary Materials for: “Induced defenses alter the strength and direction of natural selection on reproductive traits in common milkweed”

**Table S1.** Mean trait values and standard deviations, coefficients of phenotypic variation (CV_p_), and data transformations for all traits and fitness components included in the multivariate selection analysis.

| Trait | Mean | SD | $CV_p^*$ | Transformation |
| --- | --- | --- | --- | --- |
| <u>Fitness components</u> |  |  |  |  |
| Seed count | 149.97 | 307.45 | 205.00 | None |
| Pollinia removed** | 0.56 | 0.55 | 98.82 | None |
| <u>Phenotypic traits</u> |  |  |  |  |
| Floral width PC | 0.00 | 1.00 | $11.87 \pm 3.62^{***}$ | None |
| Floral length PC | 0.00 | 1.00 | $10.08 \pm 2.14^{***}$ | None |
| Date of first flower**** | 32.61 | 5.06 | 15.54 | $\ln(1 + x)$ |
| Inflorescence number | 1.94 | 1.79 | 92.10 | None |
| Inflorescence size | 47.38 | 22.65 | 47.82 | None |
| Biomass (g) | 7.79 | 7.79 | 70.79 | $\sqrt{x}$ |

\*We used untransformed data to calculate coefficients of phenotypic variation as $CV_p =$
$100(V_p^{0.5} / \mu_i)$ , where $\mu_i$ is the mean of trait $i$ .
\*\*Mean per flower
\*\*\*Mean $CV_p \pm 1$ SD of three most highly loading components (see Table S2).
\*\*\*\*Days from 1 Jun 2014.

**Table S2.** Summary of principal components analysis used to compress floral measurements into two principal components. The three variables with highest loading values on each component are highlighted in bold.

|  | Floral width (PC1) | Floral length (PC2) |
| --- | --- | --- |
| <u>Factor loadings</u> |  |  |
| Petal length | 0.16 | **0.93** |
| Petal width | **0.79** | 0.15 |
| Sepal length | 0.60 | **0.70** |
| Sepal width | **0.85** | 0.30 |
| Flower length | 0.39 | **0.90** |
| Flower width | **0.76** | 0.46 |
| Eigenvalue | 4.10 | 0.83 |
| Proportion variance explained (%) | 68 | 14 |

**Table S3.** Phenotypic correlation matrix for all measured traits*. *P*-values are below the diagonal and Pearson’s product-moment correlation r-values are above the diagonal. Significant correlations (*P* < 0.05) are in **bold.**

|  | Infl num | Infl size | Hd lgt | Hd wdt | Ptl lgt | Ptl wdt | Flwr lgt | Flwr wdt | Frst flwr | Biomass |
| --- | --- | --- | --- | --- | --- | --- | --- | --- | --- | --- |
| Infl num | — | **0.68** | **0.29** | 0.12 | **0.25** | 0.17 | **0.32** | **0.3** | **-0.31** | **0.78** |
| Infl size | **< 0.001** | — | **0.27** | 0.18 | **0.28** | 0.19 | **0.33** | **0.32** | -0.01 | **0.5** |
| Hd lgt | **0.004** | **0.008** | — | **0.38** | **0.63** | **0.43** | **0.86** | **0.49** | -0.13 | **0.46** |
| Hd wdt | 0.256 | 0.091 | **< 0.001** | — | **0.46** | **0.57** | **0.48** | **0.51** | -0.03 | **0.21** |
| Ptl lgt | **0.016** | **0.006** | **< 0.001** | **< 0.001** | — | **0.72** | **0.86** | **0.8** | -0.06 | **0.33** |
| Ptl wdt | 0.104 | 0.072 | **< 0.001** | **< 0.001** | **< 0.001** | — | **0.58** | **0.75** | -0.04 | **0.31** |
| Flwr lgt | **0.002** | **0.001** | **< 0.001** | **< 0.001** | **< 0.001** | **< 0.001** | — | **0.67** | -0.08 | **0.41** |
| Flwr wdt | **0.003** | **0.001** | **< 0.001** | **< 0.001** | **< 0.001** | **< 0.001** | **< 0.001** | — | -0.11 | **0.35** |
| Frst flwr | **0.002** | 0.920 | 0.226 | 0.740 | 0.574 | 0.683 | 0.432 | 0.305 | — | -0.11 |
| Biomass | **< 0.001** | **< 0.001** | **< 0.001** | **0.046** | **0.001** | **0.002** | **< 0.001** | **0.001** | 0.285 | — |

\**Infl num*: Inflorescence number; *Infl size*: Inflorescence size; *Hd lgt*: Hood length; *Hd wdt*:
Hood width; *Ptl lgt*: Petal length; *Ptl wdt*: Petal width; *Flwr lgt*: Flower length, *Flwr wdt*: Flower
width; *Frst flwr*: First flower

**Table S4.** Phenotypic correlation matrix for all traits* after collapsing the six floral traits into two principal components *P*-values are below the diagonal and Pearson’s *r* values are above the diagonal. Significant correlations (*P* < 0.05) are in **bold.**

|  | Infl Num | Infl Size | Frst Flwr** | Flrl Wdt** | Flrl Lgth | Biomass |
| --- | --- | --- | --- | --- | --- | --- |
| Infl Num | — | **0.68** | **-0.31** | 0.12 | **0.3** | **0.78** |
| Infl Size | **0** | — | -0.01 | 0.17 | **0.28** | **0.5** |
| Frst Flwr | **0.002** | 0.920 | — | -0.03 | -0.1 | -0.11 |
| Flrl Wdt | 0.249 | 0.096 | 0.809 | — | 0 | 0.19 |
| Flrl Lgth | **0.003** | **0.006** | 0.318 | 1 | — | **0.4** |
| Biomass | **0** | **0** | 0.285 | 0.069 | **0** | — |

\**Infl num*: Inflorescence number; *Infl size*: Inflorescence size; *Flrl lgt*: Floral length PC; *Flrl wdt*
PC: Floral width; *Frst flwr*: First flower
\*\*Principal components analysis scores

**Table S5.**
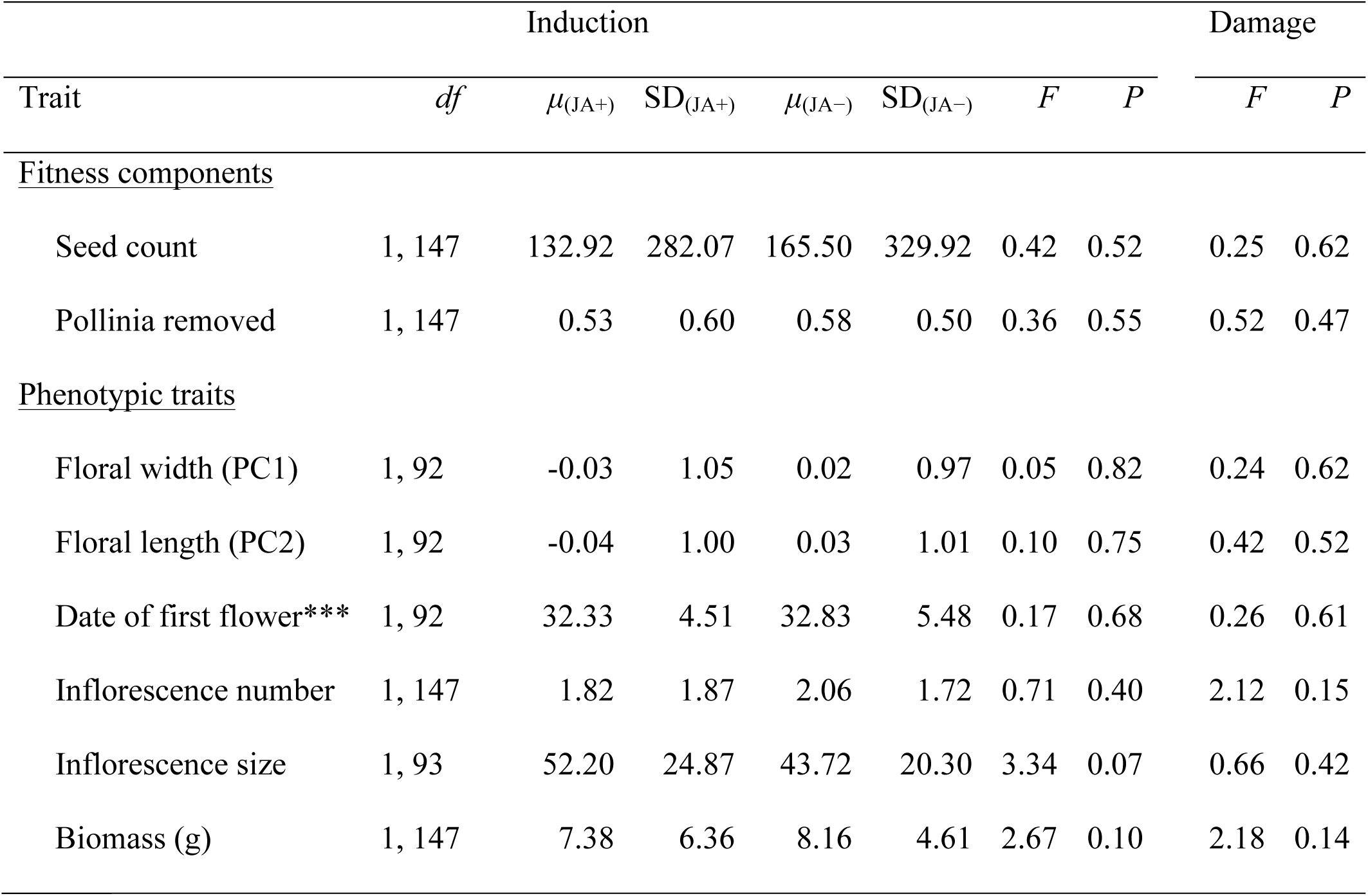
Mean values (*μ*) and standard deviations for JA-induced (JA+) and control (JA−) plants. No trait values differed between JA-induced in control plants, or across damage treatments, in univariate ANOVA with type-III SS. Mean values are not shown for the damage treatment.

**S1.2. Supplementary figures**

**Fig. S1.**
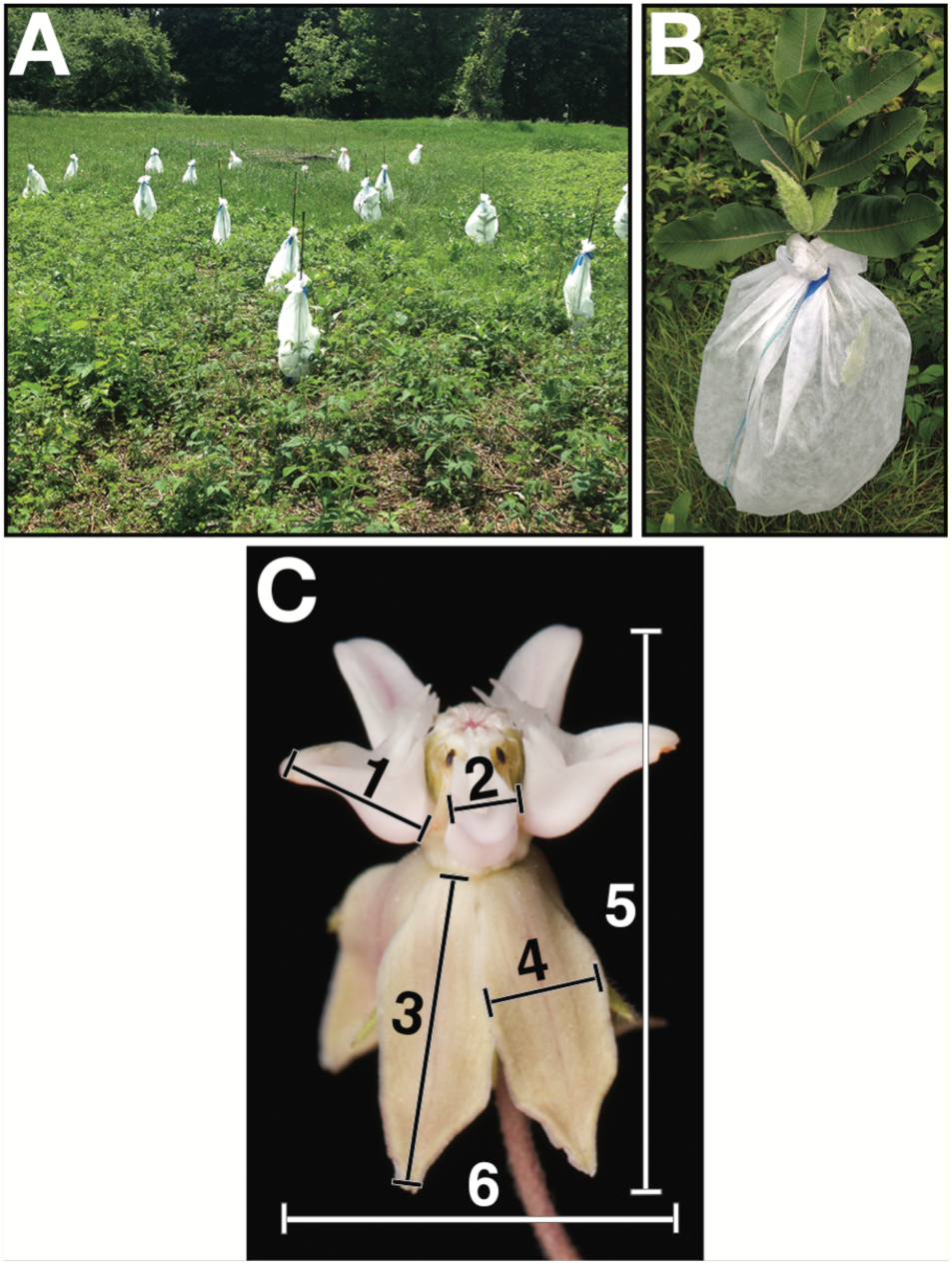
Photographs of the experiment and floral details of the common milkweed, *Asclepias syriaca*, study system. **(A)** Array of bagged milkweed plants at the beginning of the experiment. **(B)** Individual bagged milkweed plant with ripening fruit. **(C)** Photograph (credit: Ellen Woods) of an *A. syriaca* flower with flower measurements indicated (1: hood length, 2: hood width, 3: petal length, 4: petal width, 5: flower length, 6: flower width).

**Fig. S2.**
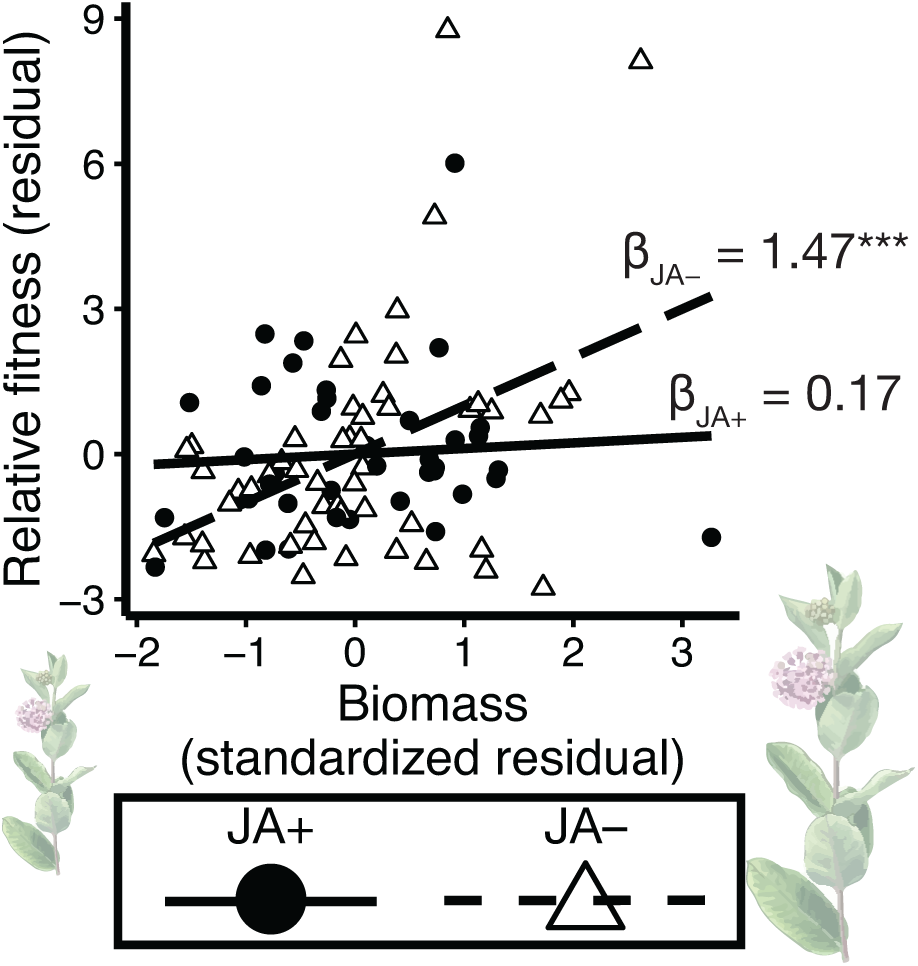
Scatterplot comparing linear selection gradients for biomass between JA-induced (*n* = 40) and control (*n* = 54) plants in the experiment. Specifically, control plants (JA−) experienced positive selection on biomass, while selection was not acting on JA-induced plants (JA+). The selection gradient values (*β*) are given adjacent to the corresponding selection gradients *(***P* < 0.001).

### Image attribution

*Asclepias* inflorescences used in Fig. 1A were modified from:

USDA-NRCS PLANTS Database / Britton, N.L., and A. Brown. 1913. An illustrated flora of the northern United States, Canada and the British Possessions. 3 vols. Charles Scribner's Sons, New York. Vol. 3: 29.

Drawings of full *A. syriaca* plants used in Fig. 1B and Fig. S2 were modified from a drawing by Zsoldos Márton, licensed under the Creative Commons Attribution-Share Alike 3.0 Unported license.

